# DeepTACT: predicting high-resolution chromatin contacts via bootstrapping deep learning

**DOI:** 10.1101/353284

**Authors:** Wenran Li, Wing Hung Wong, Rui Jiang

## Abstract

High-resolution interactions among regulatory elements are of crucial importance for the understanding of transcriptional regulation and the interpretation of disease mechanism. Hi-C technique allows the genome-wide detection of chromatin contacts. However, unless extremely deep sequencing is performed on a very large number of input cells, current Hi-C experiments do not have high enough resolution to resolve contacts among regulatory elements. Here, we develop DeepTACT, a bootstrapping deep learning model, to integrate genome sequences and chromatin accessibility data for the prediction of chromatin contacts among regulatory elements. In tests based on promoter capture Hi-C data, DeepTACT is seen to offer improved resolution over existing methods. DeepTACT analysis also identifies a class of hub promoters, which are active across cell lines, enriched in housekeeping genes, functionally related to fundamental biological processes, and capable of reflecting cell similarity. Finally, the utility of high-resolution chromatin contact information in the study of human diseases is illustrated by the association of *IFNA2* and *IFNA1* to coronary artery disease via an integrative analysis of GWAS data and high-resolution contacts inferred by DeepTACT.

## Introduction

Precise identification of physical contacts among regulatory elements is of crucial importance to not only the deciphering of transcriptional regulation, but also the understanding of the mechanism of human complex diseases. Since human variants that fall into non-coding regions are likely to be responsible for diseases [1], exploring the effects of functional variants on regulatory elements plays an indispensable role in the understanding of disease mechanism. However, most of the non-coding variants are not well annotated and not accurately linked to genes that they regulate [2], making it difficult to value the impact of these mutations. Therefore, precise identification of interactions between promoters and their regulators is urgently needed.

Over the past five years, deep neural networks have led to dramatic advances in computer vision and pattern recognition [3, 4] and have also been applied to biological problems such as the prediction of DNA accessibility and the recognition of regulatory regions and protein binding sites[5–7]. The success of previous applications of deep neural networks in biological fields inspires us to design a deep learning model to detect high-resolution chromatin contacts among regulatory elements, utilizing the advantage of deep neural networks in automatically learning meaningful feature patterns and capturing high-level context dependencies.

In this paper, we develop a bootstrapping deep learning model called DeepTACT (**Deep** neural networks for chromatin con**TACT**s prediction), to predict chromatin contacts at a resolution of individual regulatory element level using sequence features and chromatin accessibility information. DeepTACT can infer not only promoter-enhancer interactions but also promoter-promoter interactions. We show that DeepTACT improves the resolution of the high-quality promoter capture Hi-C (PCHi-C) data from multiple regulatory elements level (5-20kb) to individual regulatory element level (1kb). Besides, DeepTACT identifies a set of hub promoters, which are active across cell lines, enriched in housekeeping genes, closely related to fundamental biological processes, and capable of reflecting cell similarity. Moreover, through integrative analysis of high-resolution chromatin contacts inferred by DeepTACT and existing GWAS data, we inferred novel associations for coronary artery disease, providing a powerful way to build a fine-scale chromatin connectivity map to explore the mechanism of human diseases.

## Results

### Design of DeepTACT model and training strategy

We developed a novel bootstrapping deep learning model, named DeepTACT, to predict chromatin contacts using sequence features and epigenomic information. Specifically, the input for our predictive model consists of the sequences of two regulatory elements represented with a one-hot encoding strategy (Figure 1A) and their chromatin accessibility scores derived from DNase-seq experiments of a given cell type (Figure 1B). Based on this input, our model will compute the predictive score of whether the two regulatory elements are in physical contact. The model is a deep neural network consisting of three modules: 1) a sequence module for extracting sequence features with two convolutional neural networks (CNNs), 2) a DNase module for learning epigenomic features from chromatin accessibility scores with another two CNNs, and 3) an integration module for merging features of the former two modules and gaining higher-level context features with an attention-based recurrent neural network [8, 9] (Figure 1C).

**Figure 1:**
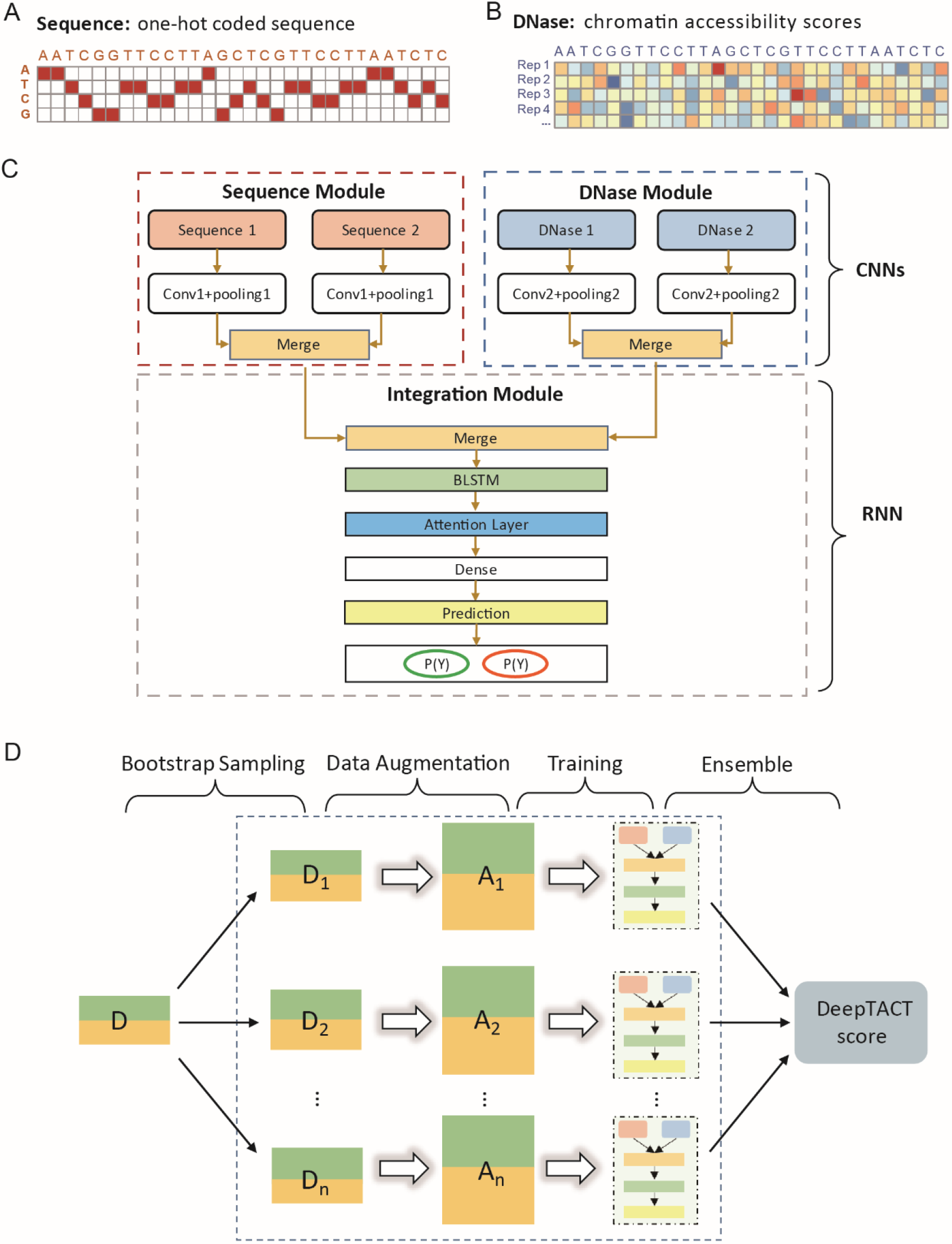
The DeepTACT method. (A) One-hot encoded sequence matrix. (B) Chromatin accessibility score matrix for replicates of a given cell line. (C) Schematic illustration of the deep neural network architecture. (D) Schematic illustration of the bootstrapping technique. See Methods for more details.

In addition, we use an ensemble technique, bootstrapping [10], to overcome the instability of the deep neural model caused by random initialization of parameters and local minimum of optimization. First, we bootstrap from an original training set to generate new subsets with the same sample size as the original set. Then, a data augmentation strategy is applied to the resulting subsets to obtain larger datasets for model training. The deep neural network of Figure 1C is trained based on each augmented subset independently, resulting in a binary classifier. The final output is an ensemble of the binary classifiers derived from different subsets (Figure 1D).

Next, we discuss the important issue of how to train the model. Since we want to make context-specific prediction, we designed a strategy to construct the training data from the chromatin contact data from that context, so the context-specific model. Suppose we have chromatin contact data in a certain context. The data consist of tens of thousands of pairs of interacting regions, where each individual region may be 5-20kb in size. While a majority of the regions contain multiple regulatory elements, a percentage (about 8.56% in the data in [20]) of the interacting pairs have the property that each region in the pair contains only one regulatory element. Our strategy is to use these pairs, which can indeed identify interacting regulatory elements, to construct the positive training cases. To the best of our knowledge, DeepTACT is the first deep learning method that considers sequence features and epigenomic information together to predict chromatin contacts among regulatory elements.

### DeepTACT accurately predicts chromatin contacts

We designed a series of experiments to systematically evaluate the ability of DeepTACT in capturing promoter-promoter interactions and promoter-enhancer interactions. Specifically, we derive training and testing samples from six sets of human PCHi-C data published in [11] as follows. If a PCHi-C interaction only contains one regulatory element (i.e. a promoter or an enhancer) on both ends, we considered this interaction as a true interaction (Figure 3A-B). Taking data processing for promoter-promoter interactions as an example, we first regarded true promoter-promoter interactions as positive samples and uniformly divided them into ten subsets: one for testing, and the others for training (Table 1). Then, we generated negative samples under the constraint that the distance between the two promoters has the same distribution as that of the positive samples. For testing sets, we generated five times more negative testing samples than positive ones. Data processing for promoter-enhancer interactions is performed in a similar way.

**Figure 2:**
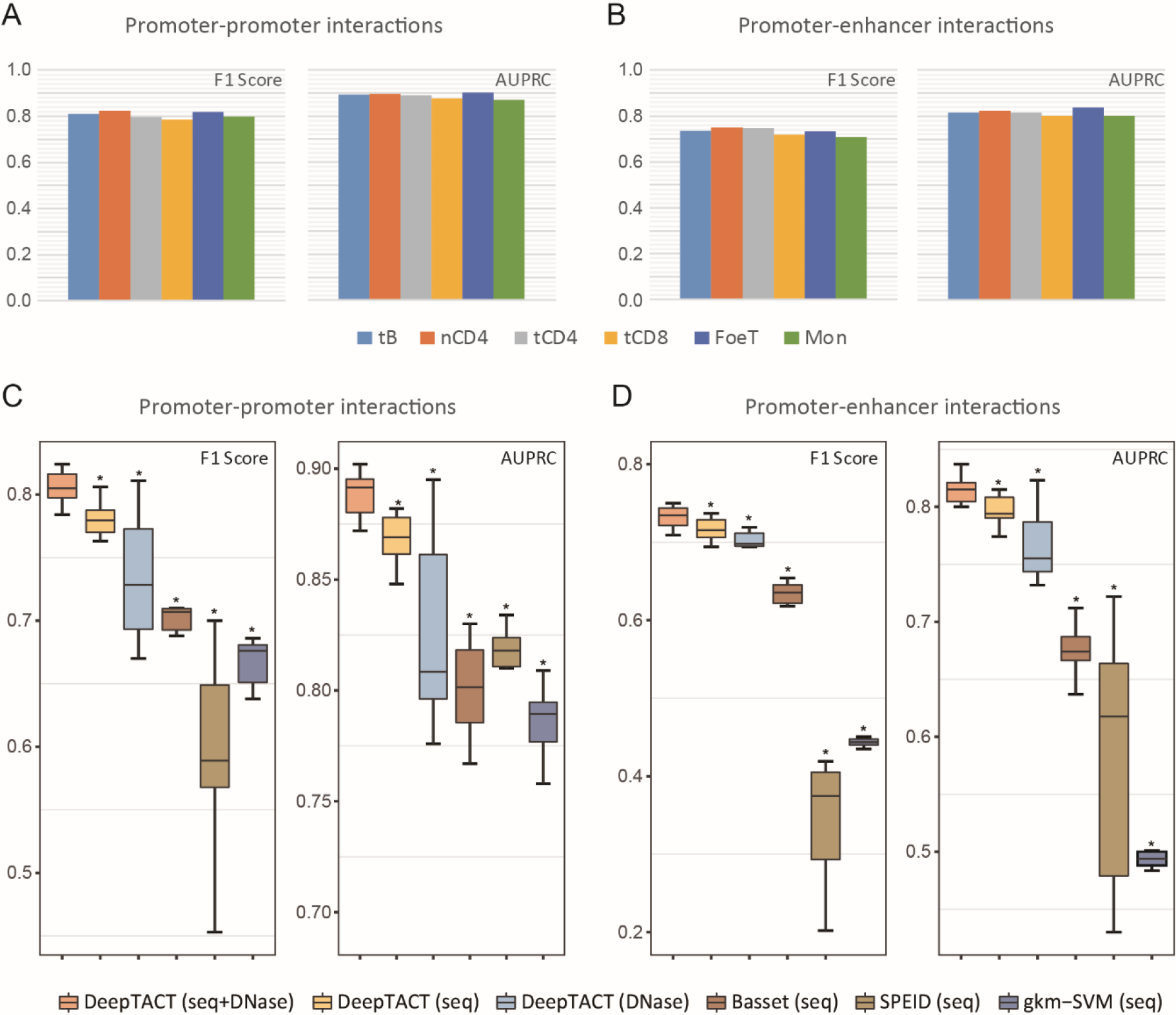
Performance evaluation of DeepTACT. (A) Prediction evaluation of promoter-promoter interactions. (B) Prediction evaluation of promoter-enhancer interactions. (C) Model comparisons of promoter-promoter interactions. (D) Model comparisons of promoter-enhancer interactions. Each box represents a distribution for six cell lines. *P* values are based on Wilcoxon signed-rank tests. * indicates *P* < 0.05.

**Figure 3:**
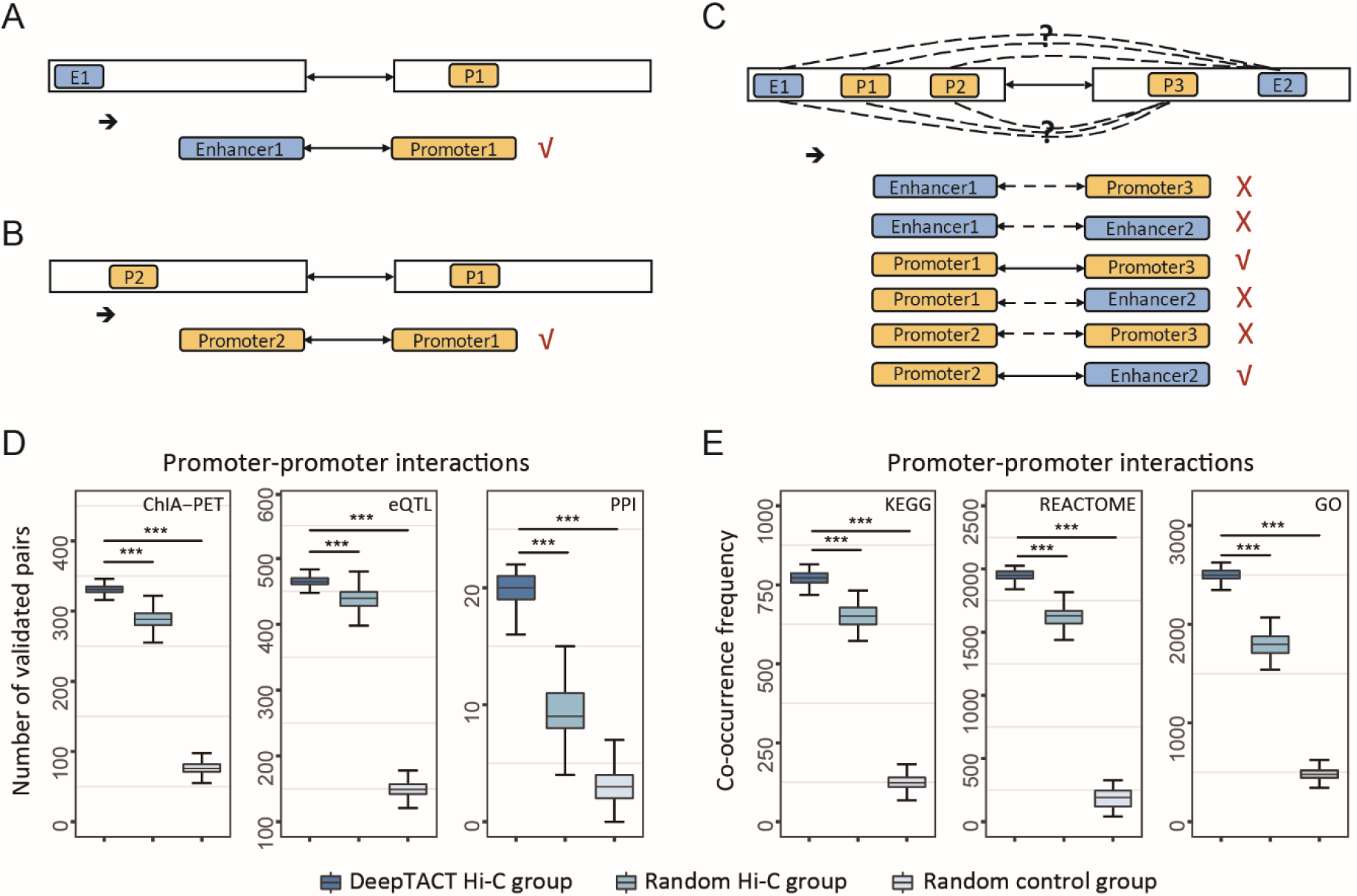
High-resolution promoter-promoter interactions. (A, B) A pair of interacting regions with only one regulatory element in each region. (C) A pair of interacting regions with one or both regions containing multiple regulatory elements, resulting in several candidate promoter-level interactions. (D) Comparisons of numbers of promoter-promoter interactions validated by ChIA-PET data, eQTLs, and PPIs. (E) Comparisons of co-occurrence frequencies of promoter-promoter interaction pairs in KEGG pathways, REACTOME pathways, and GO terms. For each type of interactions, 1,000 subgroups were sampled at the same sample size to generate a distribution. *P* values are based on one-sided Wilcoxon rank-sum tests. *** indicates *P* < 2.2 × 10^−16^.

**Table 1:** Numbers of valid positive samples for promoter-promoter interactions and promoter-enhancer interactions.

| Cell Line | Acronym | Promoter-promoter Interactions |  | Promoter-enhancer Interactions |  |
| --- | --- | --- | --- | --- | --- |
|  |  | Positive Pairs | Augmented Positive Pairs | Positive Pairs | Augmented Positive Pairs |
| **Total B cell** | tB | 8,307 | 166,140 | 9,037 | 180,740 |
| **Monocytes** | Mon | 7,016 | 140,320 | 8,353 | 167,060 |
| **Total CD4+ T cell** | tCD4 | 8,025 | 160,500 | 8,282 | 165,640 |
| **Total CD8+ T cell** | tCD8 | 7,949 | 158,980 | 8,403 | 168,060 |
| **Naive CD4+ T cell** | nCD4 | 8,612 | 172,240 | 8,995 | 179,900 |
| **Foetal thymus** | FoeT | 7,378 | 147,560 | 6,880 | 137,600 |
| **Mean** | - | 7,246 | 144,910 | 7,822 | 156,440 |

For all the six data sets in [20], DeepTACT yields F1 scores of 0.78-0.82 and AUPRCs of 0.87-0.90 for promoter-promoter interactions (Figure2A; Table 2). For promoter-enhancer interactions, DeepTACT is slightly less accurate and has F1 scores of 0.71-0.75 and AUPRCs of 0.80-0.84 (Figure2B; Table 2). We also found that DeepTACT with both sequences and chromatin accessibility data as the input outperformed modified simplified DeepTACT model with either sequences or chromatin accessibility data as the input (Figure 2C-D). This indicates that chromatin accessibility data provide complementary information to sequences in detecting cell-type specific interaction. In addition, we collected transcription factors (TFs) reported to be most closely related to each cell line [12], and we found these key TFs were captured by at least two ensemble models of DeepTACT, indicating sequence patterns learned by DeepTACT are informative. Finally, we compared DeepTACT with three other state-of-art methods: Basset [6], SPEID [13] and the gkm-SVM method [14]. On the six data sets in [20], DeepTACT acheives a mean AUPRC score of 0.89 for promoter-promoter interactions compared with 0.80 of Basset, 0.80 of SPEID and 0.78 of gkm-SVM. For promoter-enhancer interactions, DeepTACT achieves a mean AUPRC of 0.82 compared with 0.68 of Basset, 0.58 of SPEID and 0.50 of gkm-SVM (Figure 2C-D). Taken together, the above results show that DeepTACT is capable of integrating sequences and chromatin accessibility data together to identify chromatin contacts among regulatory elements.

**Table 2:** Performance of DeepTACT on imbalanced testing sets of six cell lines.

| **Cell Line** | **Promoter-promoter Interactions** |  | **Promoter-enhancer Interactions** |  |
| --- | --- | --- | --- | --- |
|  | F1 Score | AUPRC | F1 Score | AUPRC |
| **tB** | 0.811 | 0.893 | 0.737 | 0.815 |
| **Mon** | 0.799 | 0.872 | 0.709 | 0.801 |
| **tCD4** | 0.797 | 0.890 | 0.746 | 0.815 |
| **tCD8** | 0.784 | 0.877 | 0.718 | 0.800 |
| **nCD4** | 0.824 | 0.896 | 0.750 | 0.823 |
| **FoeT** | 0.818 | 0.902 | 0.732 | 0.837 |
| **Mean** | 0.806 | 0.888 | 0.732 | 0.815 |

### DeepTACT improves the resolution of promoter capture Hi-C data

Once the model has been learned, we can apply it to infer contacts between regulatory elements in situations where one or both interaction regions contain multiple regulatory elements (Figure 3A-C). We test the performance of this inference on a PCHi-C dataset of B cells at the resolution of 15kb, to check whether our model can lead to the improved inference of promoter-promoter interaction. We first collected all candidate interactions from the dataset. Then, we used the model trained in B cells to detect true interactions from all candidate pairs. To guarantee prediction precision, for each pair of interacting regions, we infer interaction only for the pair of promoters with the highest DeepTACT score.

For promoter-promoter interactions, we compared the number of interaction pairs validated by ChIA-PET, eQTLs or PPIs for inferred interactions with that for candidate interactions. Results show that inferred interactions were supported significantly more often than other interaction groups (Figure 3D). Besides, for two promoters of an interaction pair, we checked their co-occurrence in KEGG pathways [15], REACTOME pathways [16], or GO terms [17]. In all three databases, the co-occurrence frequency of inferred interactions was found to be significantly higher than those in other groups (Figure 3E).

For promoter-enhancer interactions, we compared the number of interaction pairs validated by ChIA-PET or eQTLs for inferred interactions with that for candidate interactions. We found that DeepTACT-inferred interactions were supported by the databases significantly more often than interactions of the other two groups (Figure 4A). Furthermore, we collected processed TPM values for five RNA-seq replicates in B cells. As shown in Figure 4B, regulated genes defined by DeepTACT tend to have significantly higher expression levels than those defined by PCHi-C data, indicating P-E interactions inferred by DeepTACT are more related to gene expression than those derived directly from high-quality PCHi-C data. Finally, we checked the functional enrichment of genes with distal enhancers in B cells. We found genes regulated by distal enhancers tended to be enriched for functions related to metabolic processes (Figure 4C-D), which is consistent with previous findings [18, 19]. In contrast, genes without any distal enhancer did not show significant enrichment in these processes (Figure 4E).

**Figure 4:**
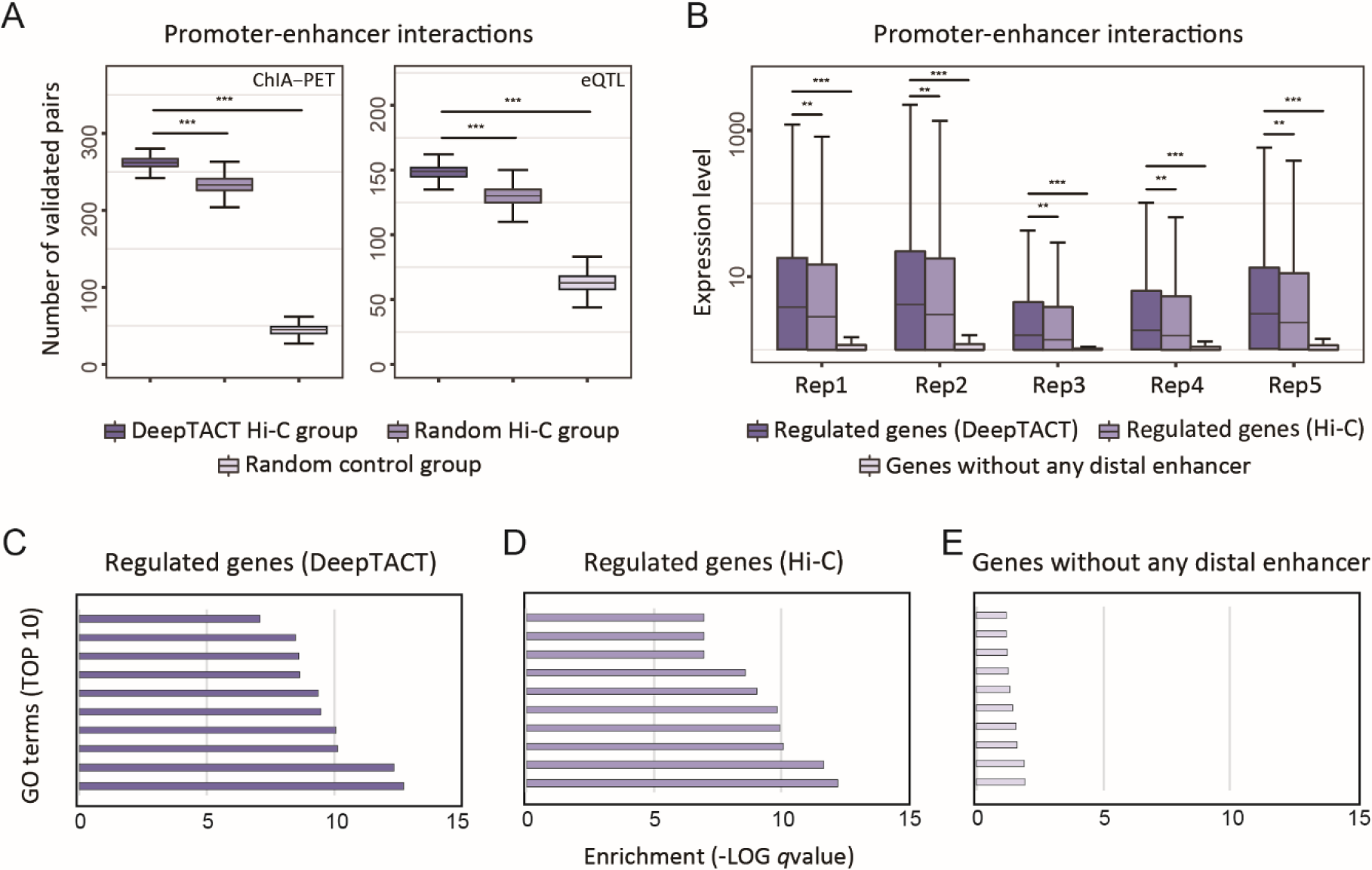
High-resolution promoter-enhancer interactions. (A) Comparisons of numbers of promoter-enhancer interactions validated by ChIA-PET data and eQTLs. (B) Comparisons of expression levels of regulated genes defined by DeepTACT interactions, regulated genes defined by random Hi-C interactions, and non-regulated genes. For each type of interactions, 1,000 subgroups were sampled at the same sample size to generate the distribution. *P* values are based on one-sided Wilcoxon rank-sum tests. *** indicates *P* < 2.2 × 10^−16^, and ** *P* < 0.005. (C) Top 10 enriched GO terms of regulated genes defined by DeepTACT interactions. (D) Top 10 enriched GO terms of regulated genes defined by random Hi-C interactions. (E) Top 10 enriched GO terms of non-regulated genes.

Collectively, these results provide evidence for the improvement of PCHi-C resolution, indicating that interactions inferred by DeepTACT have stronger biological meaning than interactions directly defined through the original PCHi-C data.

### Characterization of hub promoters defined by high-resolution interactions

We defined the top 10% promoters most frequently involved in chromatin contacts as hub promoters, yielding 1,302 hub promoters in B cells. We examined multiple genomic signals to assess the characterization of these hub promoters.

First, we collected 1,256 ChIP-seq profiles of six core histone marks (H3K4me3, H3K27ac, H3K4me1, H3K4me2, H3K9ac, and H3K9me3) from ENCODE [20]. For each histone mark, we checked the activity of hub promoters across cell lines. For comparison, we extracted promoters with the highest interaction degrees defined by candidate interactions as a control group and randomly generated non-hub promoters as another control group. As shown in Figure 5A, for histone marks enriched in transcriptionally active promoters [21–23], hub promoters defined by DeepTACT interactions were significantly more active across cell lines than the other two promoter groups. For the histone mark H3K9me3, which is related to transcriptional repression [24], hub promoters defined by DeepTACT showed the lowest activity across cell lines. Second, we checked the activity of hub promoters in 4,383 ChIP-seq profiles of 579 TFs collected from ENCODE [20]. Results show that hub promoters defined by DeepTACT interactions were significantly more active and covered more active TFs than those defined by PCHi-C data (Figure 5B). Third, we collected 3,669 housekeeping genes from [25] and detected a large fraction of overlaps between hub promoters and housekeeping genes, leading to a significantly high enrichment of hub promoters in the housekeeping genes (*P* value = 1.27 × 10^−26^, Fisher’s exact test). Fourth, we found proteins encoded by hub promoters had a significantly higher average degree compared with random protein groups (Figure 5C; *P* value = 1.30 × 10^−4^). Finally, we assessed the functional enrichment of hub promoters in GO terms [17] and REACTOME pathways [16]. Results show that hub promoters are significantly enriched in the core biological processes and pathways (Figure 5D-E).

**Figure 5:**
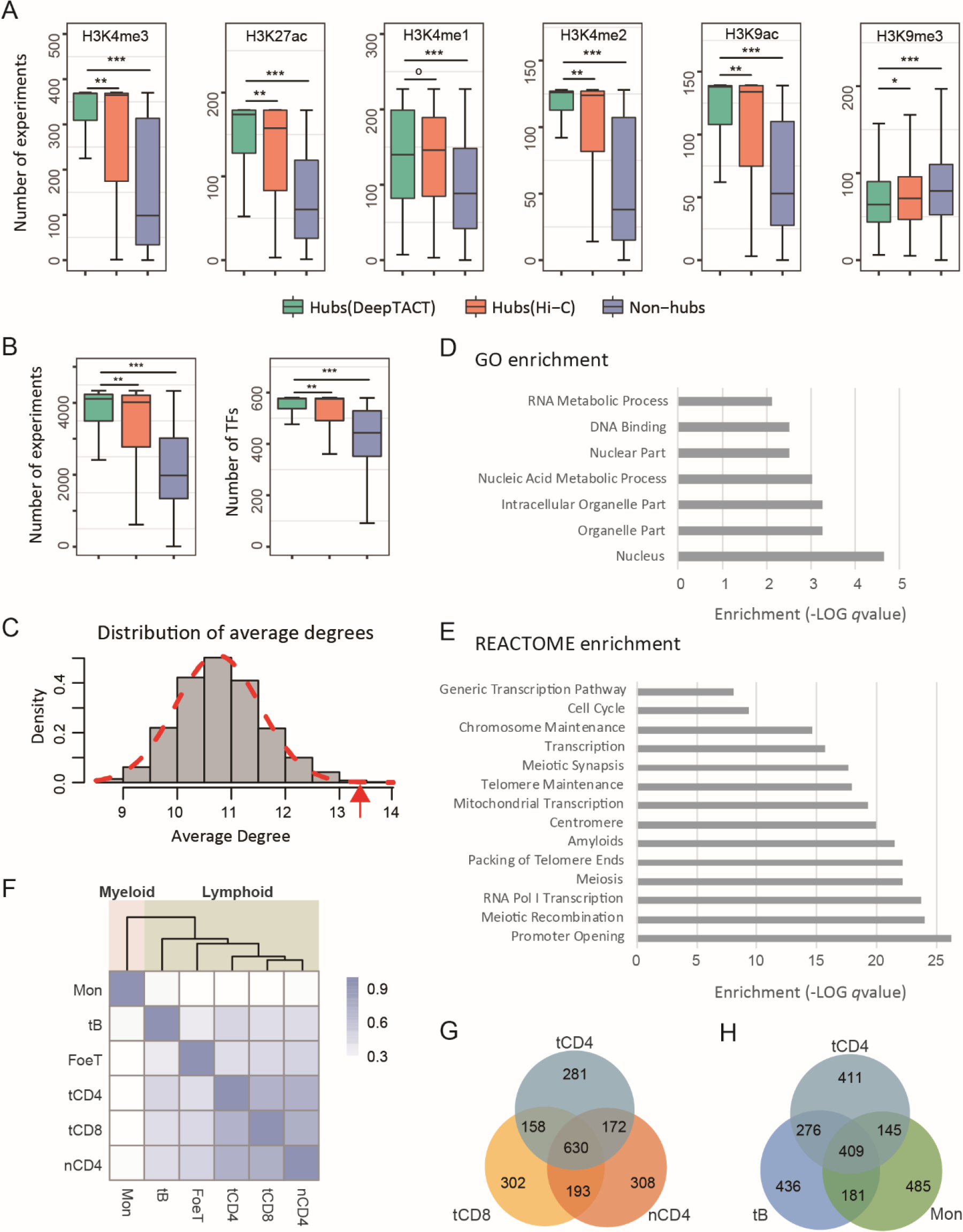
Characterization of hub promoters. (A, B) Comparisons of hub promoters defined by DeepTACT interactions, hub promoters defined by random Hi-C interactions, and non-hub promoters in terms of (A) six core histone marks and (B) TFs. The y-axis represents the number of experiments where a promoter is active. For each type of promoters, 1,000 subgroups were sampled at the same sample size to generate the distribution. *P* values are based on one-sided Wilcoxon rank-sum tests. *** indicates *P* < 2.2 × 10^−16^, ** *P* < 0.005, * *P* < 0.05, and o P > 0.05. (C) The distribution of average degrees of promoter groups. The red arrow represents the average degree of hub promoters defined by DeepTACT interactions. (D, E) Top enriched GO terms and REACTOME pathways. (F) Hierarchical clustering of the cell types according to their hub promoters. The heatmap shows the Jaccard score of each two cell types. (G) The Venn diagram of hub promoters derived from tB, tCD4, and Mon. (H) The Venn diagram of hub promoters derived from tCD4, tCD8, and nCD4.

In summary, hub promoters defined by DeepTACT interactions are characterized by distinct features. To draw a more general conclusion, we analyzed the hub promoters defined by high-resolution interactions detected in the other five cell lines and obtained similar results. We further observed the hub promoters of different cell lines and found a significantly large fraction of overlaps between each two groups of hub promoters (Figure 5F; Jaccard scores ranging from 0.291 to 0.467), supporting the point that hub promoters are fundamental across cell lines. Interestingly, we observed more overlaps among hub promoters of different types of the same cell line (i.e. tCD4, tCD8, and nCD4) than hub promoters of different cell lines (Figure 5G-H), indicating that hub promoters can reflect cell similarity. With this understanding, we applied hierarchical clustering to the six cell lines based on Jaccard scores of their hub promoters. The result reveals the lineage relationship of different cell lines, which is totally consistent with the hematopoietic tree [11] (Figure 5F).

### Identification of disease-related genes and enhancers using high-resolution interactions

We designed a computational method to identify disease-related genes and enhancers by integrating summary statistics of genome-wide association study (GWAS) data of a given disease and chromatin contacts of a cell line related to the disease. We illustrate this by an initial example. Since there have been reports of the involvement of T-cells in coronary artery disease (CAD) [26, 27], we explore the use of high-resolution interactions detected in total CD4+ T cells to identify coronary artery disease (CAD)-related promoters and enhancers. First, we collected 79,128 SNPs from the meta-analysis of GWA study including a total of 22,233 patients and 64,762 normal individuals [28]. Meanwhile, we merged all interactions inferred by DeepTACT from total CD4+ T cells, as well as positive training interactions, into a highly sparse gene-enhancer network of 22,702 nodes and 40,993 edges. After excluding genes not coding proteins and cross-chromosome interactions, we simulated a random walk process on the gene-enhancer network with *P* values of nodes derived from GWAS data as initial probabilities, yielding a steady probability score for each node. The steady scores can be served as a measurement of the association between a gene/enhancer and the disease.

As a result, we found *IFNA2* (Interferon Alpha 2), which cannot be detected based on the GWAS signal (initial *P* value = 1), ranks second by the random walk. Specifically, we found that there was no significant SNP around the TSS of *IFNA2* (the nearest signal is more than 500kb away; Figure 6A) and the initial probability of *IFNA2* was lower than other 40 genes and enhancers (Figure 6B), making it hard to associate *IFNA2* to CAD based on the GWAS signals only. Intriguingly, when integrated with the gene-enhancer network, *IFNA2* was given the second highest steady score (Figure 6C), implying that *IFNA2* may be associated with CAD. More interestingly, we found another gene-enhancer network derived from high-resolution interactions of B cells links *IFNA2* to *MLLT3*, which locates at a risk locus of leukaemia in 9p21.3 [29] and plays a key role in B-cell precursor acute lymphoblastic leukaemia [30]. Collectively, *IFNA2* is linked to genes locating at risk loci of different immune diseases in 9p21.3 by high-resolution interactions derived from different disease-related tissues, suggesting *IFNA2* is a common key autoimmune gene target. Besides *IFNA2*, we also identified two additional novel candidates for CAD, a gene *IFNA1* (Interferon Alpha 1) and an enhancer chr9:21817235-21817446 (Figure 7), which have not been reported to be related to CAD yet. Altogether, this example suggests that of joint analysis of GWA studies and the gene-enhancer network may be helpful in prioritizing disease-related genes and enhancers.

**Figure 6:**
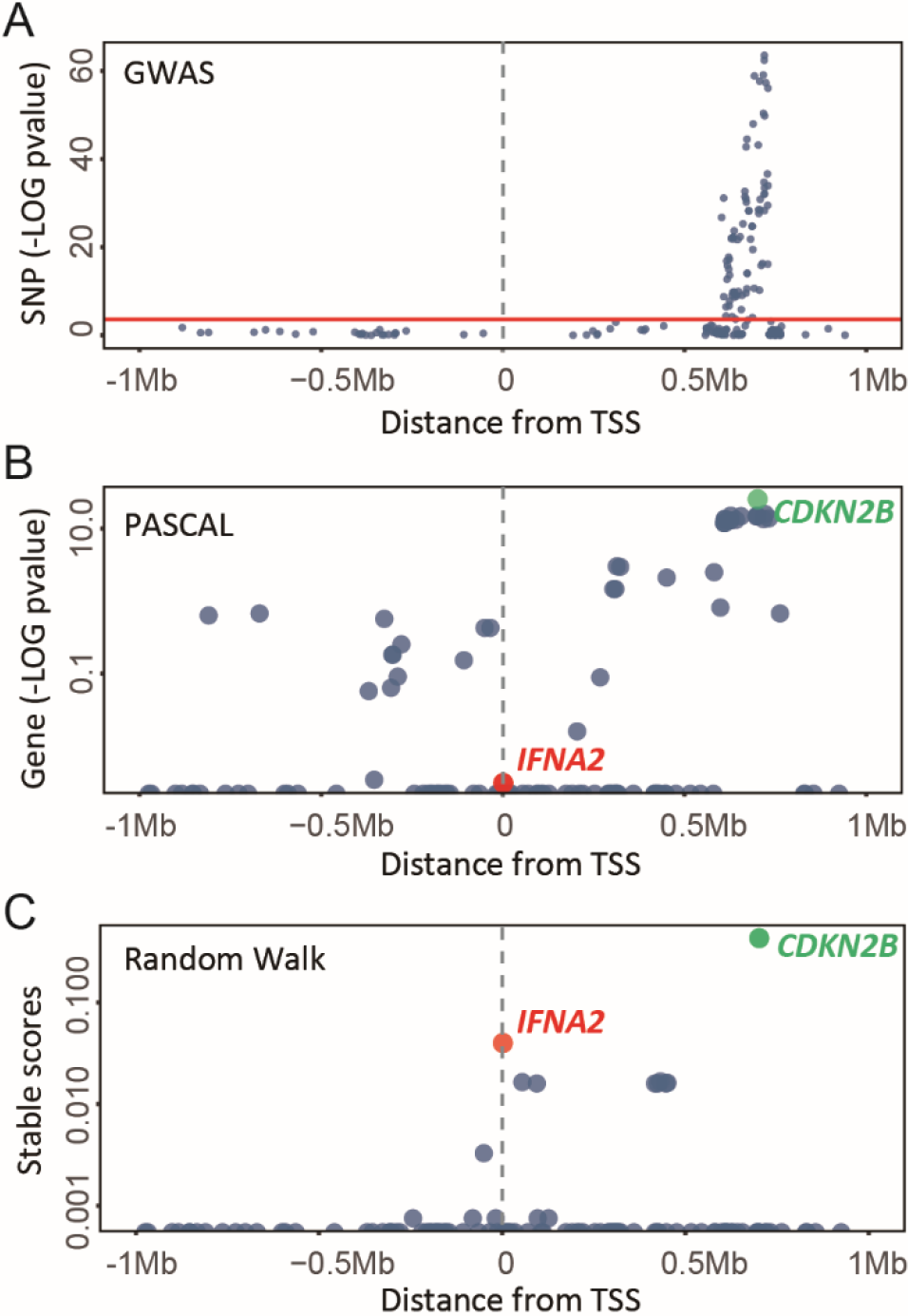
Manhattan plots of signals around *IFNA2*. (A) The Manhattan plot of GWAS SNP signals around the TSS of *IFNA2* (1Mb upstream and 1Mb downstream). The red horizontal line represents the genome-wide significance level (Bonferroni correction *P* value = 2.15 × 10^−4^). (B) The Manhattan plot of initial PASCAL *P* values of genes and enhancers around the TSS of *IFNA2.* (C) The Manhattan plot of stable scores of genes and enhancers around the TSS of *IFNA2. CDKN2B* (green) and *IFNA2* (red) were ranked first and second by the random walk.

**Figure 7:**
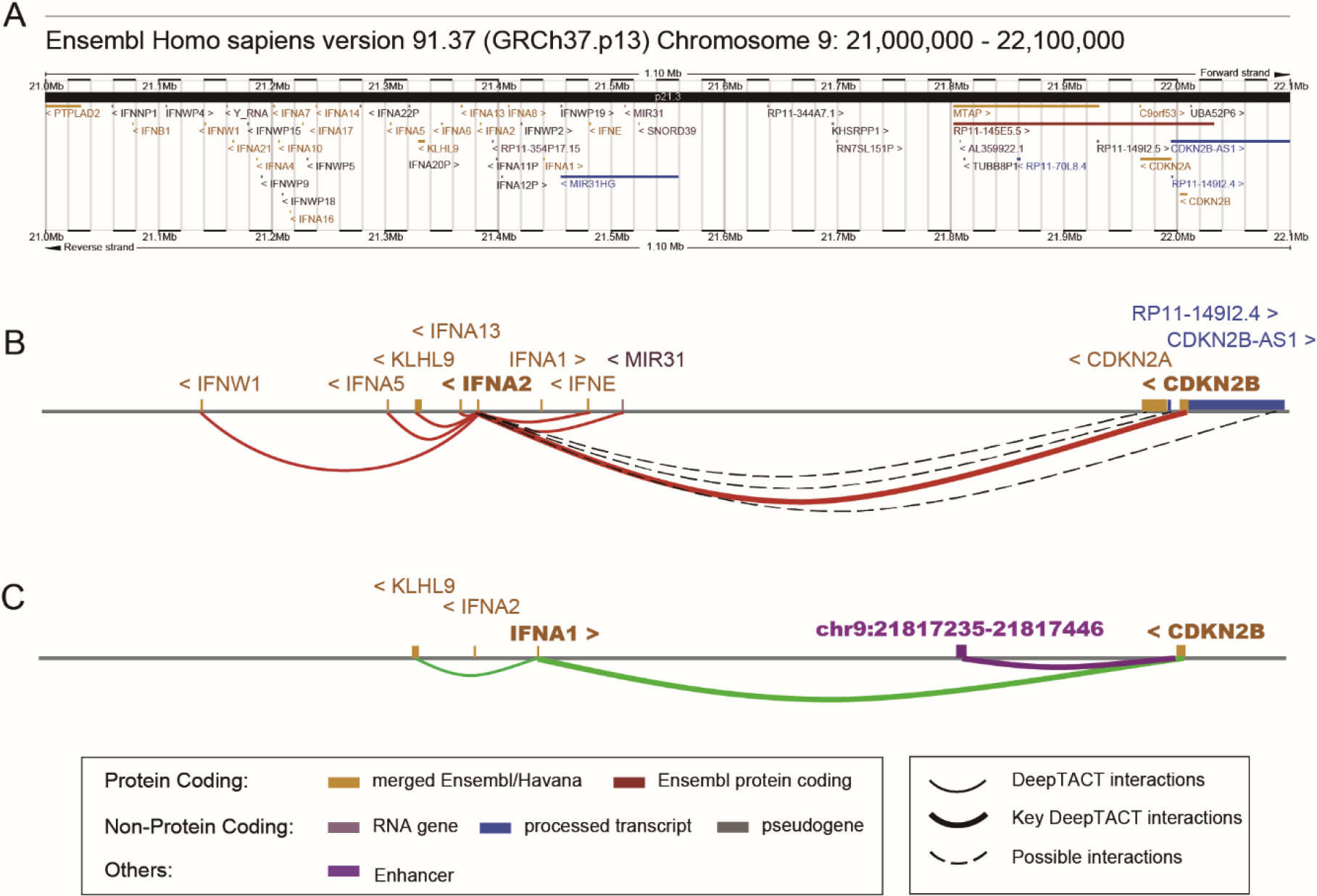
High-resolution contacts linking disease-related genes/enhancers. (A) Chromosome 9: 21,000,000 - 22,100,000 from Ensembl Homo sapiens version 91.37. (B) 3D contacts linking *IFNA2* to other genes (red curves). (C) 3D contacts linking *IFNA1* and chr9:21817235-21817446 to other genes (green curves and purple curves). High-resolution interactions are displayed with full curves, while possible interactions derived from bait-level Hi-C data are represented by grey dashed curves.

## Discussion

Several directions are worth exploring in the future. First, features learned by the model could be further explored to explain the relationship among the TFs enriched in different ends of interaction pairs. Although we have already reported a number of tissue-specific TFs learned by convolution kernels of DeepTACT from different tissues, it is hard to decipher the interactions among the TFs based on the present architecture of our model, which has only one convolution layer for each regulatory element. A specially designed strategy is needed in the future to answer this question. Second, in this work we applied the DeepTACT model to a series of promoter capture Hi-C data to improve their resolution. We expect that our model can also be applied to general Hi-C data. However, since it is more complicated to obtain training sets from general Hi-C data, the preprocessing process needs to be well designed for the further application. Third, we selected DNase-seq data to provide tissue-specific information and help achieve more powerful performance. But there are other choices to attain tissue-specific epigenomic information, such as ATAC-seq data, ChIP-seq of histone marks and TFs, and data for DNA methylation. A comparison of model performance given different types of epigenomic data as inputs will shed lights on the understanding of the relationship between different epigenomic events and the 3D contacts. Forth, high-resolution chromatin contacts connect genes to their regulators and thus help interpret the regulatory effects on the expression of target genes. This interpretation makes high-resolution interactions useful in predicting gene expression. Particularly, high-resolution chromatin contacts may create an unprecedented opportunity to handle the inherent sparsity of single-cell RNA-seq data. Finally, regulatory interactions can be used to score the influence of GWAS variants on the regulation mechanism. Given a GWAS variant falling into any end of an interaction, we can assign a score for the variant based on the difference in predicted interaction probabilities between original sequences and sequences after mutation. In this way, high-resolution interactions can offer an opportunity for the annotation and interpretation of the noncoding genome.

## Methods

### Deep neural network

The core structure of the deep neural network used in DeepCAT can be divided into three modules: a sequence module, a DNase module, and an integration module (Figure 1C). The former two modules are used to learn motif-like patterns from sequences and DNase data with separate convolutional neural networks (CNNs). In each CNN, a convolution layer is used for feature extraction, together with a rectifier operation (ReLU) to propagate positive outputs and eliminate negative outputs. Max-pooling layers are used to reduce dimensions and help extract higher-level features. In the integration module, the features learned by the above CNNs are concatenated with a merging layer, followed by a bidirectional long-short-term memory (BLSTM) layer to further learn the context features from the pooled sequence patterns. As a typical representation of recurrent neural networks (RNNs), BLSTM is widely used for its ability to capture dependencies in sequences by accessing long-range context [31]. To help the RNN pay more attention to specific sequence patterns, an attention layer is adopted in the integration module, following the BLSTM layer. The final layer of the integration module is a dense layer which is actually an array of hidden units with the ReLU activations feeding into a logistic regression unit that predicts the probability of an interaction. In addition, we adopt batch normalization layers to accelerate the training process and dropout layers to avoid overfitting.

We implemented the DeepCAT model using Keras 1.2.0 [32] on a Linux server. All experiments were carried out with 4 Nvidia K80 GPUs which significantly accelerated the training process than CPUs.

### Bootstrapping strategy

DeepCAT employs a bootstrapping strategy derived from the theory established in [10] for more stable performance (Figure 1D). We first generate *K* (*K* = 20 in this paper) new datasets of equal size as the original training set by random sampling with replacement from the training data. Then, we apply a data augmentation strategy to each new dataset *D*_*i*_, yielding an augmented dataset *A*_*i*_. After that, a deep neural network as described above is trained based on each augmented dataset independently, resulting in an ensemble of the binary classifier {*S*_*i*_}. Given the information of a sample as an input, its final prediction score is the average of the prediction scores derived from all classifiers, as

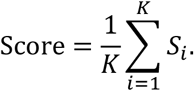

### Comparisons with other methods

We compared DeepCAT with three other methods: Basset [6], SPEID [13] and gkm-SVM [14]. Considering other methods cannot integrate sequences and DNase data together, we chose to compare the performance of all methods given only sequences as inputs and then compared the performance of DeepCAT given different inputs. Basset and gkm-SVM were not designed to deal with the interaction prediction problem, we therefore concatenated the sequences of two elements of an interaction into a single long sequence as an input, while this process was implemented by a merging layer in DeepCAT and SPEID. Briefly, we trained all methods with the same training sets and compared the performance of them with the same imbalanced testing sets to ensure the fairness of comparisons.

## Declarations

### Availability of data and material

Promoter capture Hi-C data are available in Open Science Framework (https://osf.io/u8tzp/) [11]. DNase-seq data are available through the ENCODE project [33]. Permissive enhancers are available through FANTOM5 (http://fantom.gsc.riken.jp/data/) [34]. TSS locations are available through Ensembl [35]. ChIA-PET data are downloaded from [36] and processed using a standard tool ChIA-PET2 [37]. eQTLs are available at http://genenetwork.nl/bloodeqtlbrowser [38]. RNA-seq data and ChIP-seq data for histone marks and TFs are available through the ENCODE project [33], and the accession numbers are shown in Additional file 4. KEGG pathways [15], REACTOME pathways [16], or GO terms [17] are available through MSigDB (http://software.broadinstitute.org/gsea/msigdb).

#### Funding

This research was supported by the National Natural Science Foundation of China (Nos. 61721003, 61573207 and 61175002), the NIH grants (R01HG007834 and P50HG007735), the National Basic Research Program of China (No. 2012CB316504), and the National High Technology Research and Development Program of China (No. 2012AA020401).

